# The state of preparation in performance climbing

**DOI:** 10.1101/631838

**Authors:** Simona Trifu, Antonia Ioana Trifu

**Author notes:** University of Medicine and Pharmacy “Carol Davila” Bucharest, 37 Dionisie Lupu Str., Bucharest, Romania (Simona Trifu). 21 Constructorilor Str., Bldg. J1, apt. 44, 060504 Bucharest, Romania.

## Abstract

This research has been carried out among climbing performers in Romania (a group of 60 climbers), starting from the desire to induce a state of preparation by watching a motivational short movie before performing a high difficulty route. The concept of preparation was related to the emotional impact of tonic or sensitive type (depending on the content of the movie) and the personality structure of the athletes, the conclusions drawn being in the area of optimization of performance by inducing an optimal state of preparation.

Performance climbers can have two main attitudes to impact with emotional stimuli in the competitive environment: tonic versus sensitivity. We propose the study of the correlations between the personality structure of the athletes, the emotional impact on stimulation, respectively the quality of the prepared state of state, as the active regulatory status.

The methodology included a batch of 60 climbers divided into two equal subgroups, before making a difficult route being allowed to view a movie with a tonic impact, or a sensitive impact. Personality was evaluated through five scales (Intelligence, Emotional Stability, Sensitivity, Imagination, and Perspicacity) while administering a Preparatory and Motivation Questionnaire.

People with a high level of intelligence, imagination and perspicacity can more easily create attitudes, habits and habitual contests, as well as conduct appropriate to the concrete conditions of the competitive situation, while people with low emotional stability and sensitivity are more inclined towards a sensitive, labile, sensitive approach to the competitive situation. The research implies the necessity of organizing the mental operators with the purpose of suitability to the performance poor, in accordance with the tactical training of the athlete and with the personality traits.

Emotional stimulation leads to affective participation, reception and awareness of favoring issues, stimulation of will, self-regulation of activity according to aspirations and strategies.

## 1. Introduction

### 1.1. Purpose and objectives of the research

The purpose of this research is to identify and demonstrate the existence of a relationship between affectivity and the induction of a preparation state as a specific attitude in preparation for a competition and personality type of the athletes, based on five scales of the CAQ personality questionnaire: intelligence, emotional stability, sensitivity, imagination and perspicacity.

Considering the above-mentioned purpose, research aims at achieving the following objectives:

- identifying the correlation between elevated scores of certain scales that highlight personality traits and the emotional impact of attitudinal stimulation in a visual manner (by watching an emotionally charged film), which may be tonic or sensitive.

- emphasis on the need to induce the state of preparation of climbers, as an active state with informational, decisional and energetic regulation in order to obtain a favorable competitive outcome.

### 1.2. Research hypotheses

General hypothesis: about the emotional impact of the films, there was a significant difference between two groups: Group 1 and Group 2 as will be described below.

Hypotheses derived from the general hypothesis are:

- H1: There is a negative correlation between subjects with Imagination and High Perspicacity and the emotional impact that can be seen.

- H2: There is a positive correlation between subjects with Emotional Stability and Sensitivity and emotional impact tonic

## 2. Materials and methods

### 2.1. Research variables

The present research highlights the existence of a relationship between the independent variable represented by the personality structure of the climbers participating in the study, with the five types of personality scales (intelligence, emotional stability, sensitivity, imagination and perspicacity) and the variable dependence of the emotional nature of the film. As for the emotionality of the film, the following are the indicators: emotional impact tonic on the dimension of dynamism, activism and mobilization of psychic force (supposed to be determined by the movie in which a top sportsman is successful) and emotionally sensitive impact on the dimension of empathy, raising affective and extrinsic motivation in the sense of thanking and liking others (supposedly determined by the film with the little girl who makes sacrifices for performance).

It is also noted in the H1 hypothesis that the negative correlation between 2 scales of the independent variable personality structure and an indicator of the variable depends on the emotionality of the film, as in the H2 hypothesis, there is a positive correlation between the other two scales of the independent variable personality and the second indicator of the variable depend on the emotionality of the film.

### 2.2. Participants

The study was conducted in the climbing halls in Bucharest and had athletes practicing on average three climbing exercises per week, all of them participating in competitions in the country, some of them abroad, about half of those surveyed being part of the national team over the past few years or taking part in the competition finals.

They were asked during their normal training, when they were completing a boulder route to stop trying and to watch a movie about 2 minutes long. After the movie, all the subjects completed:

- a questionnaire built specifically for this research, called the Induction Preparation Questionnaire (20 items) before the contest, together with

- CAQ Personality Inventory (Clinical Analysis Questionnaire), a short form, which has only five clinical scales (previously exposed) and which contains 59 items.

The group of climbing subjects was divided into two:

- half of them have watched a movie from the Boulder World Cup, its finals, in which a admired climber with public support, manages to top the last minute of the competition on a high difficulty route, and

- the second group watched a second movie, lasting 4.05 minutes, highlights a rhythmic gymnast at some significant moments of her childhood, moments that include both her emotional lability, sensitivity, sacrifice, the risk of giving up, and happy moments of laurel success and emotional impact on the mother.

The age of the participants is between 19 and 50 years. Two groups of participants were used: Group 1 consists of 30 participants that watched the emotionally impacting film, and the group 2 included 30 participants and watched the movie with a high emotional impact. So, the sample of this research is comprised of 60 subjects (35 female and 25 male) aged 19 to 50.

For a clear perspective on the distribution of subjects based on gender, age and emotional impact of the film, the frequency analysis was performed in the SPSS statistical processing program. Running the statistical procedures necessary to perform the frequency analyzes, we obtained the data expressing the percentage of the subjects distributed according to some indicators, as follows in table 1.

**Table 1:** Gender distribution of participants

|  |  | Frequency | Percent | Valid Percent | Cumulative Percent |
| --- | --- | --- | --- | --- | --- |
| Valid | feminine | 35 | 58.3 | 58.3 | 58.3 |
|  | masculine | 25 | 41.7 | 41.7 | 41.7 |
|  | Total | 60 | 100.0 | 100.0 | 100.0 |

**Table 2:** Age category distribution

|  |  | GENDER |  |
| --- | --- | --- | --- |
|  |  | feminine | masculine |
|  |  | Count | Count |
| Age category | 19-25 | 30 | 18 |
|  | 26-30 | 2 | 1 |
|  | 31-40 | 1 | 4 |
|  | 41-50 | 3 | 1 |

Therefore, of the total of 60 subjects tested:

- 58.3% are female,
- 41.7% are male.

A total of 60 subjects, as shown in Figure 1, is included in the following categories:

Ø 19-25 years: 48 subjects, representing a percentage of 80.00% of the total,
Ø 26-30 years: 3 subjects, representing a percentage of 5.00% of the total,
Ø 31-40 years: 5 subjects, representing a percentage of 8.33% of the total,
Ø 41-50 years: 4 subjects, representing 6.67% of the total.

Randomized sampling based on subject availability was used with regard to the sampling strategy. Choosing participants was not the age limit. In order to allow an acceptable level of test power and effect size, and for the use of parametric tests, more than 30 subjects were targeted at the time of sampling.

### 2.3. Research instruments

This research focuses on the emotional impact of a significant content and emotional film in terms of inducing the preparation state as a specific attitude to competition. Thus, the subjects watched:

- Tonic emotional impact video (a sample of 30 subjects),

- Sensitive impact video (a sample of 30 subjects).

Subsequently, all 60 subjects were asked to complete a Visually Induced Drug Testing Questionnaire (CSPIV), measured with the Likert Scale, made specifically for the current research and a Clinical Analysis Questionnaire (CAQ), from which only the following five scales have been selected:

1. Intelligence;
2. Emotional stability;
3. Sensitivity;
4. Imagination and
5. Perspicacity.

The CAQ (Clinical Analysis Questionnaire) questionnaire is an assessment tool that complements the 16PF Personality Factor Questionnaire (Cattell, 1975) and was built within the IPAT California by Krug (Forns, Maria, Amador, Juan A, Judit, Martorell, Bernardi, 1998). The CAQ questionnaire is divided into two broad groups: a group that targets the normal personality scales and traits, and a group that focuses on clinical factors analysis. In the group containing features of normal personality, the following scales are found emotional heat, intelligence, emotional stability, dominant, impulsiveness, conformism, eccentricity / extravagance, suspicion, imagination, perspicacity, insecurity, sensitivity, radicalism, self-discipline, tension.

The scales targeted in the current study are as follows:

#### Emotional stability (versus instability)

If low scores of the emotional stability scale are present, this means that there is a high anxiety state, as this feature is first affected in the pattern of anxiety, its contribution being negative. The individual level of the emotional stability factor can be considered an index of individual stress tolerance. High-score individuals are generally able to fulfill their proposed goals without difficulty. They are not easily distracted when working and are generally happy with how they work and how they live their lives. If the score of this factor is down, it is related to physical illness, suggesting an increased medical risk for chronic conditions. This scale is positively correlated with feelings of personal satisfaction and self-satisfaction.

#### Sensitivity

The association of this scale with the high scores includes: suspicion, jealousy, criticism, irritability. This feature is a normal personality trait towards paranoia, which extends to pathological dimensions. Individuals do not forget the mistakes and perceive their parents as very strict. They are concerned about the fact that others are talking back or that they may be criticized at work. Those with low scores can be considered healthy no matter how low the score is.

#### Imagination

The imagination factor does not seem to be a significant clinical factor; high-scorers are unconventional and uninterested in what happens every day and are going to forget important things, not interested in mechanical aspects. He said their princes were preoccupied with intellectual matters.

#### Agility

Subjects with high scores are sophisticated, do not express their feelings in a gentle way, being diplomatic, polite, and preferring to keep their own problems for themselves. If they have low scores, individuals are left less bound by rules and standards.

The 5 evaluated scales contain 59 items, and the Cronbach index has a value of 0.86 (table 3), indicating a high internal consistency and is located at the upper end of the estimate range tending to 1.00. The test-retest fidelity analysis was also performed, indicating the value of 0.88 when it was reapplied to a group of 30 individuals.

**Table 3:** Internal consistency index Cronbach alpha – Clinical Analysis Questionnaire

|  |  |
| --- | --- |
| Cronbach alpha coefficient | No of items |
| 0, 86 | 57 |

From the perspective of determining the degree of performance motivation, the awareness of purpose, the increase of affective and voluntary self-regulation, emotional balance, mental training, combativeness and self-confidence, after visualizing the films (emotionally emotional or emotional impact) sensitive questionnaire), a questionnaire containing 20 items - questionnaire for the visual induction of the preparation state and analyzing the individual perceptions of each individual in terms of the emotional impact expressed by each movie that the subjects viewed. It is based on a 5-step Likert Scale (1 = total agreement, 2 = agreement, 3 = indifferent, 4 = disagreement, 5 = total disagreement), indicating 5 different levels of subject views. This questionnaire was conducted by the author of present research; this has led to the need to perform an item analysis to find the value of the Cronbach’s Alpha index that shows the internal consistency of the questionnaire. Entering the values in the SPSS statistical processing program and running the procedure from the Analyze / Scale-Reliability Analysis menu resulted in an index of 0.94 (table 4), which highlights a high internal consistency.

**Table 4:** Internal consistency index Cronbach alpha – Affective Perception Scale

|  |  |
| --- | --- |
| Coefficient Cronbach alpha | No of items |
| 0, 94 | 20 |

Other tools used were:

The No. 1 movie, the tonic emotional impact, with a duration of 1 minute and 20 seconds, contains the exciting story of two people: father and son, who sit on a bench in a park, and through a heated discussion but then warm between the two, it is understood that we are given so much time to talk, but we do not have time to listen.

Movie no.2. emotionally sensitive, with a 90-minute duration, in which it is a presentation of the zodiac, in a television studio. During the 2 minutes and 90 seconds, the presenter speaks constantly, without pauses, using the same voice tone, induces a state of monotony to viewers.

### 2.4. Data collection and research design

To achieve the present study, a non-experimental design was chosen and was based on the use of psychological investigation methods such as questionnaires. Therefore, questionnaires have been applied which have allowed the evaluation of the scales of Intelligence, Perspicacity, Emotional Stability, Sensitivity, Imagination and the correlation of these scales with affectivity. The selected scales from CAQ are in direct relation to what specialty literature states as important features of climbers, traits that can be modeled by the blurring of the affectional area from the perspective of what we want by emphasizing the princeps concept of this research: increasing the competitive spirit of climbers, maintenance of the psychological freshness of athletes, successful activation under fatigue, voluntary effort. Here comes the intelligence, imagination and perspicacity from the perspective of the tactical plan, including the “dramatic” behavior, the possibility of experimenting mentally and practically of the proposed plan, the intelligent and creative self-regulation in the contest, the conclusions for the future improvement of the chances that minimize the chances competitive winnings.

The procedure underway to test hypotheses began with the presentation of informed consent, where each subject chose whether to participate in completing the questionnaires. During the pre-assessment stage, participants were instructed to enjoy relaxation and quiet in the locker room before they began their sports training of the day. The training contained information on the need to provide honest answers, presented to the subjects that there were no proper or wrong answers and consequently answered in accordance with their personality and the present status of the questionnaires. Because no time limit was specified to run the questionnaires, the subjects were free to respond to each other in their own rhythm.

Subjects were asked to provide some personal data such as age and gender.

## 3. Research results

### 3.1. Analysis of data related to descriptive statistics

Prior to conducting procedures to test the statistical significance of hypotheses, we analyzed the results of descriptive statistics tables examining the characteristics of the variables in terms of central trend, spreading and distribution.

For the implementation of parametric tests, such as distribution normality and the absence of extreme values, conditions for the normality of the distribution were imposed. Because the samples do not exceed 50 subjects, the value of the Shapiro-Wilk normality test can be analyzed.

Thus, in the normality tests of the distributions, by analyzing the two dependent variables, it is observed (in Table 5) that the normality test p (Sg.) Greater than 0.05 (emotionally sensitive impact, where p = 0.079, emotional impact tonic, p = 0.096). In this case the normal distribution is confirmed.

**Table 5:** Normality tests of distributions

| The emotional level of the subjects as a result of viewing the videos | Kolmogorov-Smirnov |  |  | Shapiro-Wilk |  |  |
| --- | --- | --- | --- | --- | --- | --- |
|  | Statistic | df | Sg. | Statistic | df | Sg. |
| Emotional-tonic Impact | .137 | 30 | .157 | .916 | 30 | .079 |
| Sensitive emotional impact | .143 | 30 | .123 | .941 | 30 | .096 |

**Table 6:**
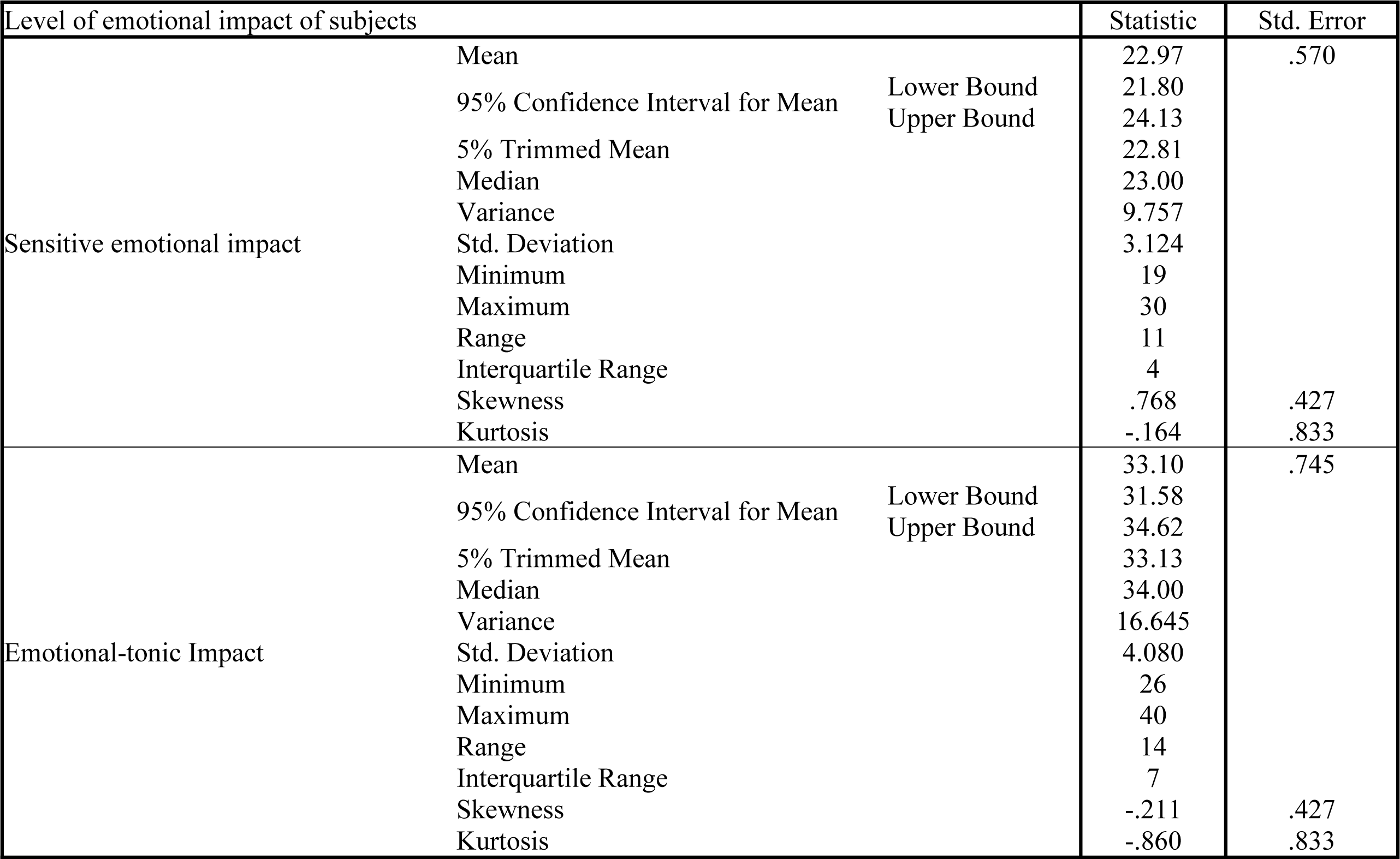
Descriptive statistics of level of emotional impact of subjects

In the following table (table 7), descriptive statistical data for a measurement scale called Emotional Stability is provided, where the average value is 7.27 and the standard deviation is 2.37. The scale is within the range of confidence limits of 6.65 and 7.88. From the analysis of the skewness symmetry index, a value of −0.45 is observed, and the value of the numerical indicator of flattening, kurtosis, is −0.78. It is noted that the data do not exceed + 1 / −1, which shows the distribution symmetry in terms of the skewness index and the kurtosis numeric indicator.

**Table 7:** Descriptive statistics for emotional stability of subjects

| Emotional Stability Level of the subjects |  |  | Statistic | Std. Error |
| --- | --- | --- | --- | --- |
| Emotional Stability | Mean |  | 7.27 | .307 |
|  | 95% Confidence Interval for Mean | Lower Bound | 6.65 |  |
|  |  | Upper Bound | 7.88 |  |
|  | 5% Trimmed Mean |  | 7.37 |  |
|  | Median |  | 8.00 |  |
|  | Variance |  | 5.656 |  |
|  | Std. Deviation |  | 2.378 |  |
|  | Minimum |  | 2 |  |
|  | Maximum |  | 10 |  |
|  | Range |  | 8 |  |
|  | Interquartile Range |  | 4 |  |
|  | Skewness |  | -.454 | .309 |
|  | Kurtosis |  | -.784 | .608 |

Table 8 provides descriptive statistics on the measurement scale, called Sensitivity. It can be seen from the table that the average has a value of 6.85 and that of the standard deviation is 2.16. The confidence limits of the variable have values of 6.29 and 7.41. The value of the skewness symmetry index is −0.37 and that of the kurtosis flattening index is −0.21. It also notes that these values do not exceed + 1 / −1, which signals distribution symmetry.

**Table 8:** Descriptive statistics for sensitivity

| Level of sensitivity for the subjects |  |  | Statistic | Std. Error |
| --- | --- | --- | --- | --- |
| Sensitivity | Mean |  | 6.85 | .280 |
|  | 95% Confidence Interval for Mean | Lower Bound | 6.29 |  |
|  |  | Upper Bound | 7.41 |  |
|  | 5% Trimmed Mean |  | 6.94 |  |
|  | Median |  | 7.00 |  |
|  | Variance |  | 4.706 |  |
|  | Std. Deviation |  | 2.169 |  |
|  | Minimum |  | 1 |  |
|  | Maximum |  | 10 |  |
|  | Range |  | 9 |  |
|  | Interquartile Range |  | 2 |  |
|  | Skewness |  | -.378 | .309 |
|  | Kurtosis |  | -.219 | .608 |

Table 9 shows descriptive indices for the measurement scale representing the level of Imagination of the subjects. Note the average value of 7.50, but also the standard deviation value, which is 2.52. The scale called Imagination has its value between the confidence limits of 6.85 and 8.15. It is also noted that the values of the skewness symmetry index are −0.97 and the kurtosis flattening index is 0.34. The value of the skewness symmetry index and that of the kurtosis flattening index do not exceed + 1 / −1, which signals the distribution symmetry.

**Table 9:**
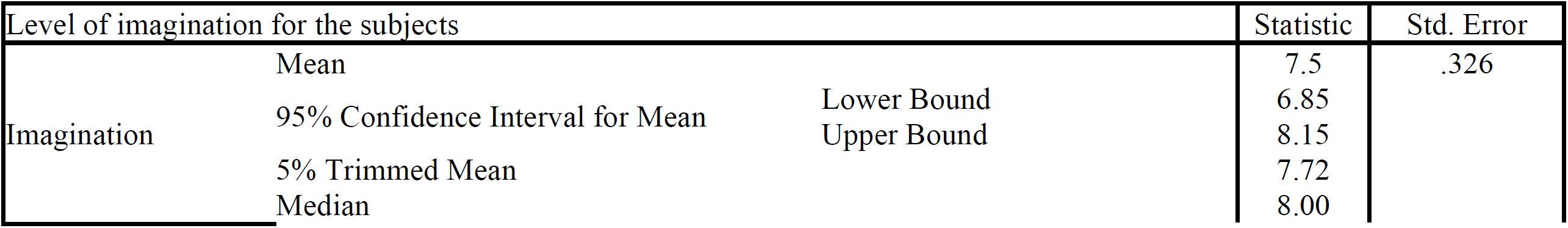
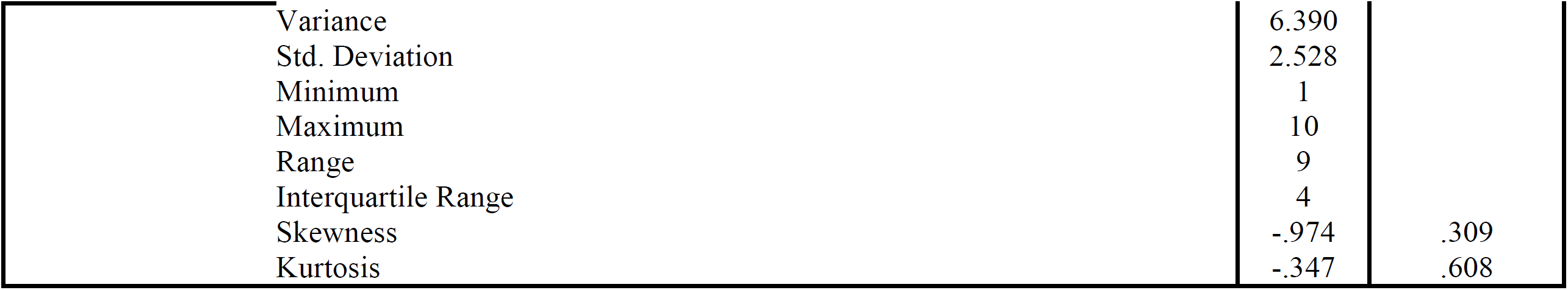
Descriptive statistics for imagination

| Level of imagination for the subjects |  |  | Statistic | Std. Error |
| --- | --- | --- | --- | --- |
| Imagination | Mean |  | 7.5 | .326 |
|  | 95% Confidence Interval for Mean | Lower Bound | 6.85 |  |
|  |  | Upper Bound | 8.15 |  |
|  | 5% Trimmed Mean |  | 7.72 |  |
|  | Median |  | 8.00 |  |

Table 9: Descriptive statistics for imagination
|  |  |  |
| --- | --- | --- |
| Variance | 6.390 |  |
| Std. Deviation | 2.528 |  |
| Minimum | 1 |  |
| Maximum | 10 |  |
| Range | 9 |  |
| Interquartile Range | 4 |  |
| Skewness | -.974 | .309 |
| Kurtosis | -.347 | .608 |

In the last table, (table 10) there is a descriptive statistic of the measure of the level of Perspicacity of the subjects who participated in the test. So, the average value is 6.77 and the standard deviation is 2.62. The Perspective Level of the subjects ranges between the confidence limits of 6.09 and 7.44. The value of the skewness symmetry index is −0.68, and the kurtosis flattening index is − 0.35.

**Table 10:** Descriptive statistics for perspicacity

| Level of perspicacity for the subjects |  |  | Statistic | Std. Error |
| --- | --- | --- | --- | --- |
| Perspicacity | Mean |  | 6.77 | .339 |
|  | 95% Confidence Interval for Mean | Lower Bound | 6.09 |  |
|  |  | Upper Bound | 7.44 |  |
|  | 5% Trimmed Mean |  | 6.91 |  |
|  | Median |  | 7.00 |  |
|  | Variance |  | 6.894 |  |
|  | Std. Deviation |  | 2.626 |  |
|  | Minimum |  | 1 |  |
|  | Maximum |  | 10 |  |
|  | Range |  | 9 |  |
|  | Interquartile Range |  | 4 |  |
|  | Skewness |  | -.684 | .309 |
|  | Kurtosis |  | -.358 | .608 |

### 3.2. Analysis of inferential statistics data

The data obtained from the questionnaire completion by the participating individuals were analyzed, as previously noted, in order to test the statistical significance of the hypotheses, the values being entered in the statistical data base analysis SPSS. Before this information was scored in the SPSS, both questionnaires went through the scoring stage, so the final scores were reached. Thus, the final scores for the scales: Emotional Stability, Sensitivity, Imagination and Perspicacity, and for the Visual Induction Questionnaire were obtained.

To test the general hypothesis, the t test for two independent samples was applied, because it was intended to show a significant difference between the averages obtained for the same variable, the emotional impact (affectivity). It was measured in two groups, the difference being the type of movie each group viewed.

From Table 11, data on homogeneity of distribution can be seen, Levene test result indicating a = 0.062, with a value greater than p = 0.05, indicating the homogeneity of the dispersions of the two groups. Next, we note the value of t = 10,80, associated with the propensity of 0,0005, df = 58, under the homogeneity of the variant. The confidence interval (95%) for the difference between the averages (mdif = 10,13 is between the lower limit (8,25) and the upper limit (12,01), and an increased precision of the average estimate is expressed. of the association between the dependent and the independent variable, the omega-square was calculated and the value ω2 = 89 was obtained, which, according to Cohen, demonstrates that there is a strong association between the variable dependence and the independent one.

**Table 11:**
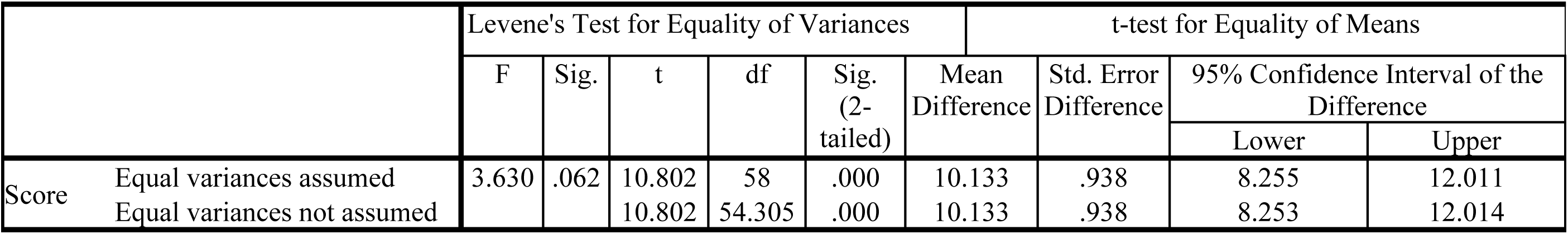
Independent Samples t-test

Testing the H1 hypothesis was done through the Pearson correlation test because it was intended to highlight a correlation between the independent variable, which is called the personality structure expressed by the Imagination and Perspicacity scales and the dependent variable, called the emotionality of the film expressed by the type of film to which they have the subjects participated before completing the questionnaire. Table 12 shows that from the scale correlation called Imagination and the sensitive emotional impact variable, there is a negative correlation between Imagination and emotionally sensitive impact, since r = −0,37 and p = 0,043, where p <0,05 and resulting negative correlation between Perspicacity and emotionally sensitive, but insignificant, because r = −0.165, p = 0.38. As both correlations are negative, H1 is confirmed.

**Table 12:** r-test of linear correlation Pearson

|  |  | Imagination | Perspicacity | Low emotional impact |
| --- | --- | --- | --- | --- |
| Imagination | Pearson Correlation | 1 | .100 | -.372 |
|  | Sg. (2-tailed) |  | .449 | .043 |
|  | N | 60 | 60 | 30 |
| Perspicacity | Pearson Correlation | .100 | 1 | -.165 |
|  | Sg. (2-tailed) | .449 |  | .384 |
|  | N | 60 | 60 | 30 |
| Senzitive emotional impact | Pearson Correlation | -.372 | -.165 | 1 |
|  | Sg. (2-tailed) | .043 | .384 |  |
|  | N | 30 | 30 | 30 |

Testing the H2 hypothesis was also accomplished through the Pearson linear correlation test, aiming to highlight a positive correlation between the scales in the personality inventory called Emotional Stability and Sensitivity and emotional impact tonic. Their bilateral testing has a significance threshold of α = 0.05. Table 13 shows that from the correlation of the scales called Emotional Stability and Sensitivity and the emotional impact, a positive correlation is observed, since between Emotional Stability and emotional impact tonic, r = 0.194, p = 0.057, where p <0.05, Here we see a positive correlation, and from the perspective of the correlation between Sensitivity and tonic emotional impact, r = 0.168, p = 0.041, where p <0.05, there is a positive correlation between the two. In conclusion, H2 is confirmed.

**Table 13:** r-test of linear correlation Pearson

|  |  | Emotional stability | Sensitivity | Tonic emotional impact |
| --- | --- | --- | --- | --- |
| Emotional stability | Pearson Correlation | 1 | .185 | .194 |
|  | Sg. (2-tailed) |  | .156 | .057 |
|  | N | 60 | 60 | 30 |
| Sensitivity | Pearson Correlation | .185 | 1 | .168 |
|  | Sg. (2-tailed) | .156 |  | .041 |
|  | N | 60 | 60 | 30 |
| Tonic emotional impact | Pearson Correlation | .194 | .168 | 1 |
|  | Sg. (2-tailed) | .057 | .041 |  |
|  | N | 30 | 30 | 30 |

## 4. Psychological interpretation of results

Following the elaboration of the current research on the link between the personality structure, concerning the scales of Emotional Stability, Sensitivity, Perspicacity, Intelligence and Imagination and the effect of the visual induction of the preparation state, it was noted, by analyzing the statistical data obtained following the application of the parametric tests, with a high level of intelligence, imagination and perspicacity can more easily create attitudes, habits and habits of competition, as well as conduct appropriate to the concrete conditions of the competitive situation, whereas people with low emotional stability and sensitivity are more inclined towards a sensitive, labile, sensitive approach to the competitive situation.

The correlation to the Pearson correlation test between the Emotional Stability, Sensitivity and Emotional Impact / Preparative Status was positive because the sample of 60 participants predominated in those with low scores at these scales and those with a low score in the first from the perspective of the Emotional Stability Scale, they generally fail to be able to fulfill their proposed goals, attention can easily be distracted when working; Secondly, from the perspective of those with a low level of the Sensitivity scale, it includes safety, but also neglect, prefers loud music and the use of force instead of reason to achieve certain results.

The second hypothesis that was confirmed was the negative correlation between Imagination, Perspicacity, and the emotionally sensitive impact of subjects as a result of watching movie no. 2, which stimulates the empathic, human emotional component, sensitivity, sacrifice, empathy and the theme of failure. From a psychological point of view, it can be stated that the analysis resulted in a negative correlation, because the sample of 60 participants predominated subjects with a high level of Perspicacity, namely Intelligence and Imagination. Subjects with a high level of Imagination are unconventional and typically uninterested in what is happening day by day; they can forget the trivial things, they are not interested in the mechanical aspect, and they can say that their parents have been or are concerned with intellectual issues, while people with a high level of Perspectivity are sophisticated, do not express their feelings easily, are diplomats, polite, preferring to keep their problems for themselves.

Thus, by extrapolating, the diachronic aspect of the preparation for the contest has in the center the psychic factor, which is made up of the athlete’s attitude system, a system that develops and progresses gradually, finally forming the specific psychic state for the contest. From a psychological point of view, the constant need for new tracking strategies makes climbers feel happy, and also provides a drop in anxious tendencies.

Terms such as motivation, goal orientation, role consciousness, self-confidence, combativeness, route approach plan, competition plan are the core of training for a competition. As a result, pre-competitive preparation requires the emotional state of mind that largely determines the mental and practical function of the performance at that time. It is a mental organization of the psychic system, with the purpose of suitability for the poor performance and in accordance with: the technical tactical training of the athlete and his personality traits.

### 4.1. Discussion

The present paper had in mind both the knowledge of the psychological mechanisms used in the induction of the preparation state as a necessary attitude for the contest, as well as the relation between the personality structure of the athletes and the emotional impact that a certain visual stimulus in symbolic significance may have upon them with reference to the personal competition past. It was previously observed in the application of the statistical test t for two independent samples because it was intended to reveal a significant difference between the averages obtained for the same variable, the emotional impact. The statistical results have the relevant value t = 10,80, associated with the probability p = 0,0005, df = 58, under the homogeneity of the variant. From the point of view of the effect size index, the value of ω2 = 89 was obtained, indicating, according to Cohen’s table, that there is a strong association between the variables. Also, the statistical data revealed the positive correlation between the Emotional Stability, Sensitivity and Tonic Emotional Impacts, resulting from the Pearson correlation test, r = 0.194, where p = 0.057, associated with p <0.05, shows that there is a correlation positive between the Emotional Stability Scale and the emotional impact of the participants, and r = 0.168 and the value of p = 0.041 shows a positive correlation between the Sensitivity Scale and the tonic emotional impact of the participants.

From the point of view of the correlation between the Imagination - Emotional Sensitive Impact scale and the Perspicacity Scale - Sensitive Impact, we found that according to the Pearson correlation test, r = −0.37 and p = 0.043 respectively r = − 0.165, where p = 0.38.

## 5. Conclusions with practical applicability in climbing

The concept of preparation requires the athlete to function in an intelligent manner, whether directed or independent (depending on circumstances), not only by learned schemes but also in a creative way. The state of preparation is the preamble of the introduction to the competitive behavioral mood, which is recommended as a competitor model that represents a synthesis of the most effective behaviors considered from the point of view of all psychic laws: intelligence, imagination, perspicacity, critical thinking, rapid analysis and synthesis alternatives, high prosexic status, both in a voluntary and spontaneous manner, increased ability to concentrate, stability and selectivity of attention.

Always the state of preparation for achieving the performance peak is directly related to the personality of the athlete, his individuality and attributes, the characteristics in which the person concerned has undergone physical training, and the general psychological training.

The preparation condition implies:

- providing optimum level of motivation to the highest level, along with the formation of the feeling of triumph in relation to the competition in fact;

- clear awareness of the proposed goal, together with the awareness of the role of the athlete and the aspiration to achieve maximum performance;

- achieving a good emotional balance in the ability to master emotions and the ability to adapt to stress;

- ability to use mental training;

- realizing the lucidity of peer and thought in critical and unexpected situations;

- adaptability of the motor reactions in relation to the requirements of the route;

- activating feelings of boldness, perseverance, courage, initiative;

- maintaining confidence in their own strengths and ability to achieve the proposed performance;

- increasing combativity and a desire for success;

- maintaining psychic prospecting by using invisible training (autogenous training);

- developing psychic resistance and immunity to frustration over unpleasant difficulties and disturbing factors.

The preparation state builds on previous direct contest experiences and can be conscious and conscious without the need to be analyzed in all its aspects. Sport mastery is defined by the ability to instantly reach the preparation state in an intuitive way. It involves the sustained volunteer effort of updating skills and abilities, sometimes postponement, suspension of immediate tendencies and reactions, for greater satisfaction in the future. It also implies rigor with regards to aspirations and level of visitation, so that the proposed goal is not far beyond the technical and emotional performance level of the stage.

The preparation state also involves the construction of the competition plan, including possible variants for difficult moments (steps). A certain level of mental tension, that constructive nervousness is required as a symptom of activation, but in a highly controlled manner. In case of over-excitement, the person knows what to do to calm himself. In the case of loss of self-confidence, the athlete possesses intrinsic internal mechanisms, so that his or her confidence can be recaptured. In the preparation state, the focus is on the future, the athlete has the self-consciousness of the measure in which he will succeed in the contest, he is able to master the unexpected situations and the unfamiliar does not affect his performances.

Through this study, we have tried to show to what extent certain personality traits, such as: emotional stability, intelligence, imagination and perspicacity, influence the competition approach and the formation of specific attitudes in the cognitive, affective and volitional area to provide the athlete maximum capability of adaptation. We started from the idea that visual stimulation by watching short films with emotional content enhances the athlete’s ability to anticipate, ideally, before approaching the path, activates the motives and diminishes pre-competitive anxiety. It also stimulates the athlete’s engagement size, in line with their performance.

Intelligence, imagination and perspicacity, as personality traits, are directly related to the anticipatory process, looking at a detailed and operationalized mental plan, as well as the design of conduct. The mental project is transformed into a concrete way and involves: accurate orientation of thoughts, mental programming, fixation of motivation, aspirations and expectations, autosuggestion of voluntary combativity and assembly.

From the perspective of our study, scales of emotional stability, intelligence and perspicacity imply a tonic emotional response to watching the movie.

The imagination and sensitivity scales are consistent with the size of emotivity, sensitivity, responsiveness to memories, and the reduction of the critical filter in terms of upgrades burdened by both past successes and failures. These two personality traits correlate with sensitive emotional involvement and are typical of athletes who use their motivation, desire, hope, and the rest of the affectionate volunteer area.

Thus, the personality traits in the area of intelligence and perspicacity refer to the ability of the athlete to focus his attention on the objective and driving actions, intensity of concentration, stability and resistance to attention to external and internal disturbances, to the state of vigilance in waiting and follow. These features involve observation, analysis and rapid synthesis of situations, lucidity in critical moments, originality, suppleness, fluidity of thought, anticipation capacity, optimal rational decision-making.

The area of imagination involves the ability to represent route strategies, inner speech (self-commands, encouragement, control of proposed conduct), motoric memory, topography, and action schemes.

The features that define emotional stability imply emotional maturity, adaptive balance, self-mastery, resistance, optimism, personal sense of self, self-acceptance, correct self-assessment, positive tone, control of fear, trance and psychosomatic disorders, agitation, control of aggression, enhancing combativity, eliminating low affective tone.

The state of preparation, as a specific attitude of success in the competition, can be amplified by concurrent visual and auditory stimulation through short films with an impaired impact that aim at: fighting the fear of failure and suffering, increasing self-reliance, eliminating negative thoughts, and amplifying positive. Emotional stimulation leads to affective participation, reception and awareness of favored aspects, empathy, stimulation of volunteer effort, doubled by self-regulation of activity according to aspirations and strategies. The end is the goal orientation, ambition to achieve goals, initiative, maintaining discipline and tactics, self-reliance, combativeness, perseverance and ambition.

Symbolically, the state of preparation means the unity / congruence between words and deeds, the safety / reliability of the functioning of the actual psychism, liveliness, energy, activism, creativity in approaching the paths.

